# Detection of anomalous high frequency events in human intracranial EEG

**DOI:** 10.1101/782912

**Authors:** Krit Charupanit, Indranil Sen-Gupta, Jack J Lin, Beth A Lopour

## Abstract

**Objective:** High-frequency oscillations (HFOs) are a promising biomarker for the epileptogenic zone. However, no physiological definition of an HFO has been established, so detection relies on the empirical definition of an HFO derived from visual observation. This can bias estimates of HFO features such as amplitude and duration, thereby hindering their utility as biomarkers. Therefore, we set out to develop an algorithm that detects high frequency events in the intracranial EEG that stand out from the background and does not require assumptions about event amplitude or morphology.

**Method:** We propose the anomaly detection algorithm (ADA), which integrates several unsupervised machine learning techniques to identify segments of data that are distinct from the background. We apply ADA and a standard HFO detector using a root-mean-square amplitude threshold to intracranial EEG from 11 patients undergoing evaluation for epilepsy surgery. The rate, amplitude, and duration of the detected events and the percent overlap between the two detectors are compared.

**Result:** In the seizure onset zone (SOZ), ADA detected a subset of conventional HFOs. In non-SOZ channels, ADA detected at least twice as many events as the standard approach, including some conventional HFOs; however, ADA also identified many low and intermediate amplitude events missed by the standard amplitude-based method. The rate of ADA events was similar across all channels; however, the amplitude of ADA events was significantly higher in SOZ channels, and the threshold between SOZ and non-SOZ channels was relatively consistent across patients.

**Significance:** ADA does not require human supervision, parameter optimization, or prior assumptions about event shape, amplitude, or duration. It provides an unbiased estimate of HFO features, and our results suggest that amplitude may differentiate SOZ and non-SOZ channels. Further studies will examine the utility of HFO amplitude as a biomarker for epilepsy surgical outcome.

## 1 Introduction

Twenty to forty percent of patients with epilepsy will not achieve seizure freedom using medication, and this may lead them to consider surgery as a treatment option.^1^ The surgical procedure often relies on localization of the seizure onset zone (SOZ) using intracranial electroencephalography (iEEG) to guide resection. Recent studies have shown that high frequency oscillations (HFOs) occur more frequently in the SOZ,^2–9^ and the surgical removal of brain regions with high incidences of HFOs has been correlated to a higher likelihood of seizure freedom after surgery.^2,3,10–13^ These results suggest that HFOs may be a valuable marker for localization of epileptogenic tissue during surgical planning. Moreover, HFOs occur in interictal periods, so their use may enable clinicians to shorten the duration of invasive monitoring.

HFOs are empirically defined as spontaneous electrographic patterns consisting of at least four cycles of an oscillation in the frequency range of 80–500 Hz, with a high amplitude that is distinguishable from the background.^14–16^ Because these are transient events, detection of HFOs is a critical step in the localization procedure. The gold standard for detection is visual identification,^8,16–18^ but automated detectors are increasingly being implemented to save time and improve reliability and reproducibility.^19,20^ Automatic detection algorithms generally consist of the same basic steps: they identify a period of increased high frequency energy (measured with root-mean-square (RMS) amplitude,^17,21,22^ amplitude of rectified filtered data,^23,24^ line length,^25,26^ Hilbert envelope,^3,4,27^ or as a peak in the time-frequency decomposition^28^), then verify that the event exceeds a minimum duration or a minimum number of oscillations. Many detection algorithms include additional steps to merge consecutive events and reject false positives.

Despite the large number of automated algorithms that are currently available, there are two challenges of HFO detection that have not yet been addressed. First, both visual and automated detection rely on the empirical definition of an HFO derived from visual observation.^14^ There is currently no clear physiological definition that can be used to guide the selection of detection parameters such as amplitude, duration, and number of cycles, as studies have shown significant overlap between pathological and physiological HFOs.^26,28–31^ However, the optimization of such parameters is critical to the accuracy of the detector.^17,23,24^ This is directly related to the second challenge: existing detection methods require complex optimization procedures. These algorithms typically contain at least 3-5 interrelated parameters, and the detection accuracy is highest when the parameters are optimized for individual subjects.^24,32^ As a result, these algorithms do not easily generalize to new datasets, and the time and effort needed for validation and optimization is a barrier to implementation in a clinical setting.

Here, we describe how to implement a new algorithm for detection of transient high frequency events in iEEG data that addresses these two challenges. Rather than identifying events with specific features, our anomaly detection algorithm (ADA) detects any events that stand out from the background, regardless of amplitude or morphology. This could include conventional HFOs, oscillations similar to HFOs but with lower amplitude, oscillations with irregular amplitude profiles, artifacts (if they are present in the data), and other unique patterns. ADA integrates several machine learning techniques, including anomaly detection, time series pattern matching, clustering, and classification. It is fully automated and does not require parameter optimization or prior assumptions about the shape, amplitude, or duration of the events. We first present our algorithm, then demonstrate its use on human iEEG data and compare the detection results to those of a standard HFO detection algorithm. We hypothesize that ADA will enable unbiased estimation of HFO properties, which has the potential to lead to the development of more accurate biomarkers of the epileptogenic zone.

## 2 Methods

### 2.1 Patients and recordings

Intracranial EEG recordings were collected from 36 adult patients between April 2015 and December 2017 at University of California, Irvine, Medical Center. All patients had medically refractory epilepsy and were undergoing electrode implantation to localize the SOZ for possible surgical resection. For inclusion in our analysis, the recordings had to fulfill the following criteria: 1) the SOZ was clearly localized to one or more iEEG channels by experienced neurophysiologists (JJL and IS) based on seizures that occurred during the monitoring period; 2) electrode locations were confirmed using co-registered pre-implantation and post-implantation structural T1-weighted magnetic resonance imaging scans; 3) a minimum recording duration of six hours with no seizures, collected overnight while the patient was resting, was available; and 4) a minimum sampling frequency of 2 kHz was used. In total, recordings from 11 patients (five female, 38.2±16.9 years old) met these criteria. The recordings were 20 to 90 hours in duration with a 2 kHz (one subject) or 5 kHz (ten subjects) sampling rate and contained data from a total of 1148 electrodes (104.3±33.1 electrodes per patient). Electrodes that could not be clearly localized to the gray matter, electrodes with continuous electrographic artifact, and electrodes within the regions of immediate seizure spread outside of the SOZ were excluded from the analysis. We analyzed the remaining 55 SOZ electrodes (average of 5.0±3.0 channels per patient), and 122 channels outside the SOZ (which we term nSOZ; average of 11.1±6.3 channels per patient). We then selected multiple three-minute segments of iEEG for each patient using the following rules. The segments were clipped from overnight iEEG records between 11 PM to 6 AM. Each segment was at least one hour away from a seizure onset time, and we ensured that segments from the same patient were separated by at least 15 minutes. This resulted in the selection of approximately three segments per hour from each subject. In total, 118 segments were analyzed (mean of 10.7±2.8 segments per patient or 32.1±8.4 minutes per patient, with a range of 7 to 17 segments per patient).

### 2.2 Anomaly Detection Algorithm

The novel algorithm described here aims to separate anomalous high frequency events from the baseline background signal without human supervision or assumptions about the appearance of the events. The procedure consists of three parts: 1) preprocessing, 2) constructing a distance matrix, and 3) clustering and classification (Figure 1). All data analysis procedures were implemented in MATLAB 2018b (MathWorks, MA) using custom-written code, and the code for the anomaly detection algorithm will be provided as supplementary material.

**Figure 1:**
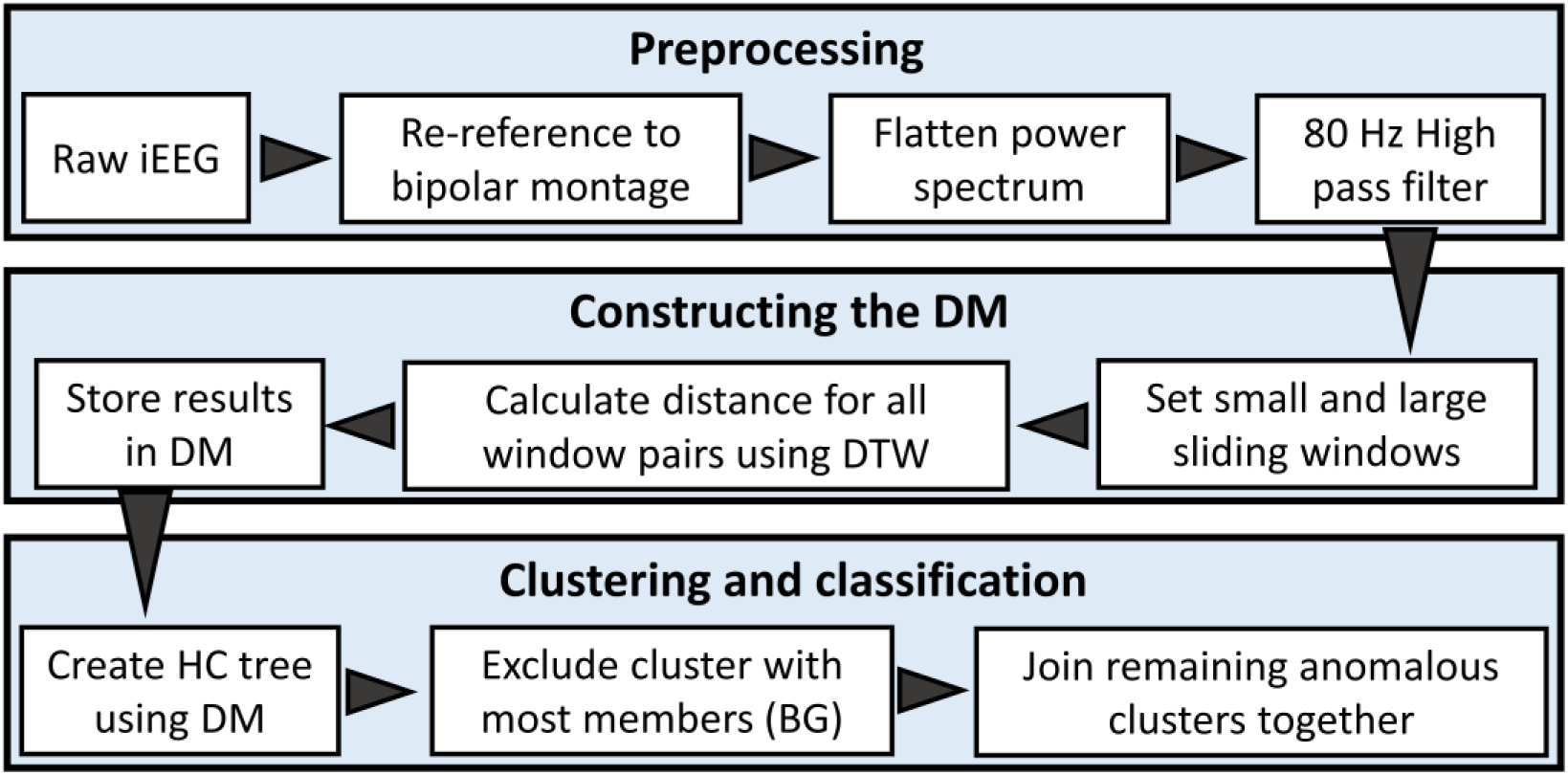
Data-flow diagram for ADA. The algorithm consists of three main parts (large shaded boxes): preprocessing, constructing the DM, and clustering and classification. Each small white box represents a major processing step within each main part, and the arrowheads show the flow of the algorithm. Abbreviations are DM: distance matrix, DTW: dynamic time warping, HC: hierarchical cluster, BG: background cluster.

#### Preprocessing

The iEEG was re-referenced to a bipolar montage via subtraction of adjacent electrodes, and the resulting signals were modified using the Simple Diff method.^33,34^ This method flattened the power spectrum in the frequency domain by enhancing the energy of the high frequency activity and suppressing the low frequency components. The flattening process started with application of the fast Fourier transform (FFT) to the data. Then the FFT power spectrum was multiplied by a constant scalar factor of 1-cos(2*π*f/fs), where f was the frequency and fs was the sampling rate. The final modified iEEG signal in the time domain was obtained by applying an inverse FFT to the modified flattened power spectrum. Each modified iEEG segment was then filtered using an 80 Hz high-pass finite impulse response digital filter. The data were filtered in the forward and reverse directions to avoid phase distortion using the filtfilt function in MATLAB.

#### Constructing the distance matrix

To construct the distance matrix for one segment of data from one channel, we first selected two window sizes: a 1.5 ms small window and 50 ms large sliding window. The small window was used to downsample the iEEG, which was necessary to reduce processing time; within each small window, the representative amplitude of the iEEG was calculated as the average amplitude of all data points. The large 50 ms sliding window was used with 50% overlap for event detection. Each large window consisted of 33 small windows, which was long enough to contain a typical HFO event.

We then measured the distance between the iEEG time series in all pairs of large windows, using the MATLAB function dtw to calculate dynamic time warping (DTW).^35,36^ DTW measures the similarity between two temporal sequences while being robust to phase differences. Then the distance matrix (DM) was created by assembling these calculated distances into a nonnegative, square, two-dimensional symmetric matrix with elements corresponding to the pairwise distances.

#### Clustering and classification

The upper triangular elements of each DM were then converted into vector form for the linkage function used to create a hierarchical cluster tree. The unweighted average distance linkage method was applied as an algorithm for computing the distance between clusters. For clustering, a maximum of seven clusters was used, although comparable results were achieved with a maximum number of clusters ranging from seven to thirteen. The cluster containing the highest number of members (where each member was one 50 ms window of iEEG data from the associated electrode) was designated as the background cluster, and the remaining six clusters were merged together and defined to be the anomaly group. Finally, for each iEEG electrode, any overlapping 50 ms windows within the anomaly group were merged into single events.

### 2.3 RMS detector

The RMS detector has become one of the most widely used automated detectors in publications related to HFOs.^2,6,34,37,38^ It is based on the moving average RMS amplitude of the 100 to 500 Hz bandpass filtered signal (finite impulse response filter, rolloff -33 dB/octave). The parameters for the RMS detector used in our study matched the original publication.^21^ We calculated the RMS amplitude using a three ms sliding window and the RMS threshold was defined as five standard deviations (SD) above the mean RMS value of the entire length of the signal. Segments of iEEG were considered to be HFO candidates when the RMS amplitude exceeded the threshold for at least six ms. Consecutive candidate events less than ten ms apart were joined together as a single event. Finally, the candidate events were accepted as HFOs if at least six rectified peaks exceeded a second threshold, defined as three SDs above the mean of the rectified filtered signal.

### 2.4 Characteristics of Detected Events

Hereafter, we will refer to events identified by the RMS detector as conventional HFOs (cHFO). Events identified via ADA, which do not have specific thresholds for amplitude or number of oscillations, will be referred to as anomalous high frequency activity (aHFA).

We compared the results of ADA and RMS detection by analyzing the shapes and characteristics of three groups of events: events detected only by ADA, only by RMS, and both by ADA and RMS, which we will refer to as ADA-only, RMS-only, and RMS+ADA, respectively. The shape of the events in each group was compared using the amplitude envelope from the Hilbert transform (median, 95^th^ percentile, and maximum). We also measured four characteristics of the aHFA and cHFO: 1) rate, defined as the average number of events per minute per channel, 2) amplitude, defined as the average value of the upper amplitude envelope over the duration of the event, 3) duration, and 4) coefficient of variation (CV). The CV is a statistical measure of dispersion of data around the mean, represented by the ratio of the SD to the mean. We used the CV to measure the consistency of the event characteristics across all segments of data. All characteristics of aHFA and cHFO were compared between SOZ and nSOZ channels. We employed a Wilcoxon rank sum test to determine whether the event characteristics were significantly different, and the significance for all analyses was set at p<0.05, except where adjusted to correct for multiple comparisons.

## 3 Results

### 3.1 Incidence and morphology of aHFA and cHFO

Across all iEEG electrodes, a total of 598 SOZ and 1360 nSOZ three-minute epochs from 11 patients were analyzed (Figure 2A). Overall, 21294 cHFOs (14008 in SOZ and 7286 in nSOZ) were detected using the RMS detector, and 22311 aHFA (6335 in SOZ and 15976 in nSOZ) were detected using ADA (Figure 2B).

**Figure 2:**
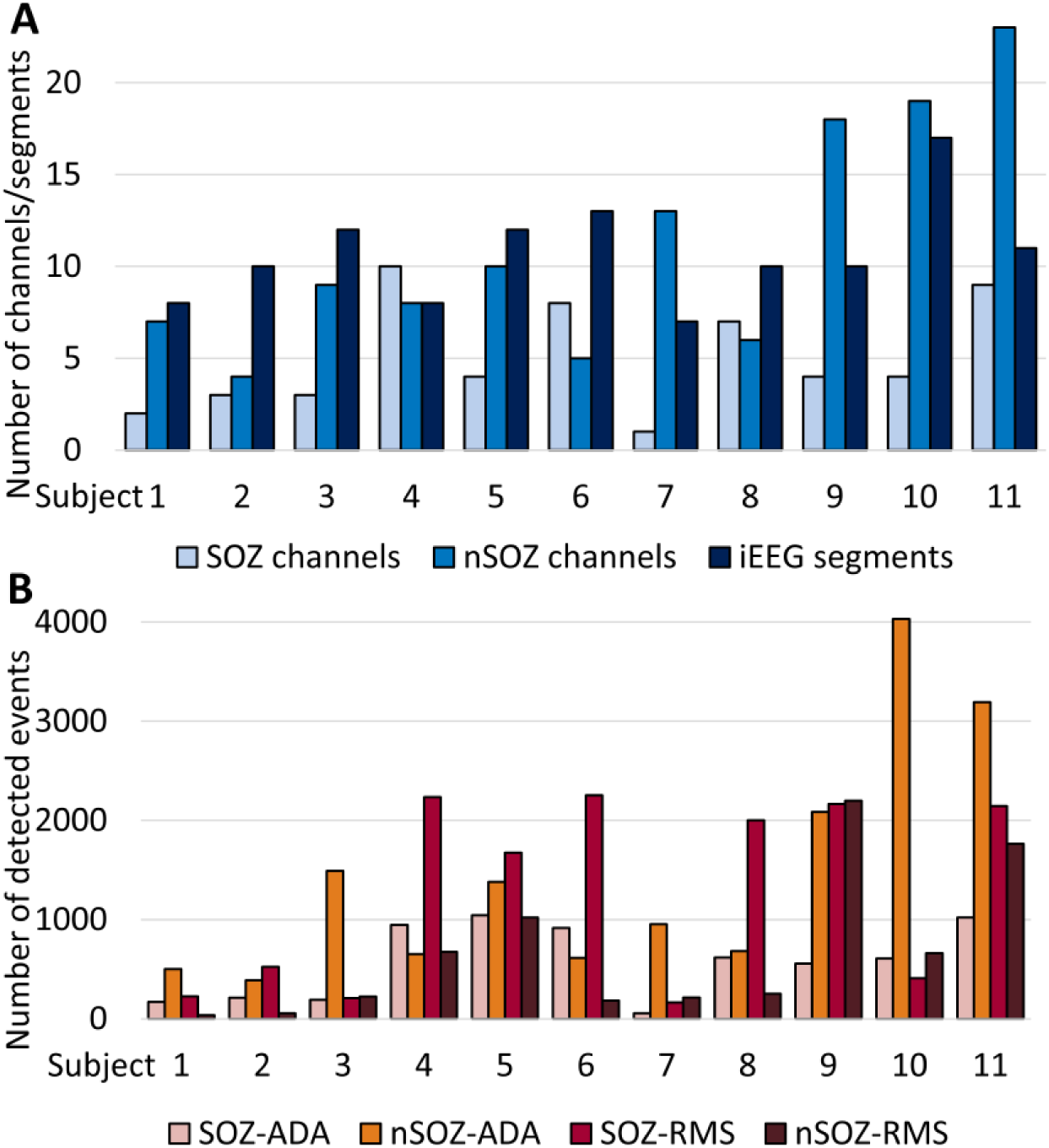
A) Total numbers of SOZ and nSOZ channels and number of three-minute segments of iEEG data analyzed for each individual patient. B) Total number of detected events from each individual patient divided into SOZ and nSOZ channels.

The number of identified events in RMS+ADA, ADA-only, and RMS-only groups varied greatly between SOZ and nSOZ (Figure 3). In every subject, a subset of events was detected by both the ADA and RMS detectors. In the SOZ, most events were RMS-only (55.5%), while 39.1% of events were RMS+ADA and 5.4% were ADA-only. In five subjects, less than 5% of events in the SOZ were ADA-only, indicating that the events detected by ADA were a subset of those identified by the RMS detector. In contrast, most events in the nSOZ channels were ADA-only (65.9%), while 10.9% were RMS+ADA and 23.2% were RMS-only.

**Figure 3:**
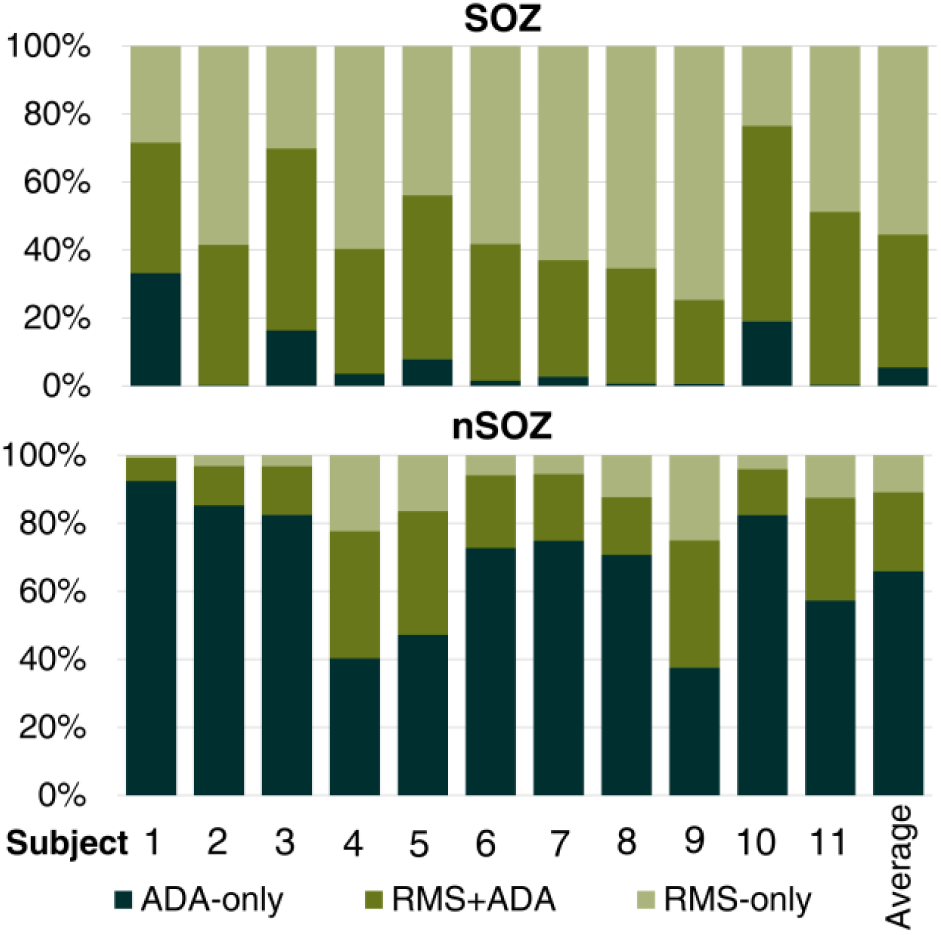
Percentages of the detected events in the ADA-only, RMS+ADA, and RMS-only groups for each subject for SOZ channels (top) and nSOZ channels (bottom).

To compare the morphology of events, we plotted the median, 95^th^ percentile, and max amplitude envelopes for each group of events (Figure 4). All groups of events exhibited amplitude profiles that reached a peak at the center of the event and tapered off at the edges, consistent with the traditional definition of an HFO. The amplitude of events in the RMS-only group was higher than those in the ADA-only group, suggesting that ADA may have considered a number of high amplitude events to be non-unique. The low-amplitude events in ADA-only were likely missed by the RMS detection scheme due to the application of a strict amplitude detection threshold. This result is consistent with the data in Figure 3; because ADA detects low-amplitude events, aHFA represent a larger percentage of events in nSOZ channels, where high amplitude events are infrequent.

**Figure 4:**
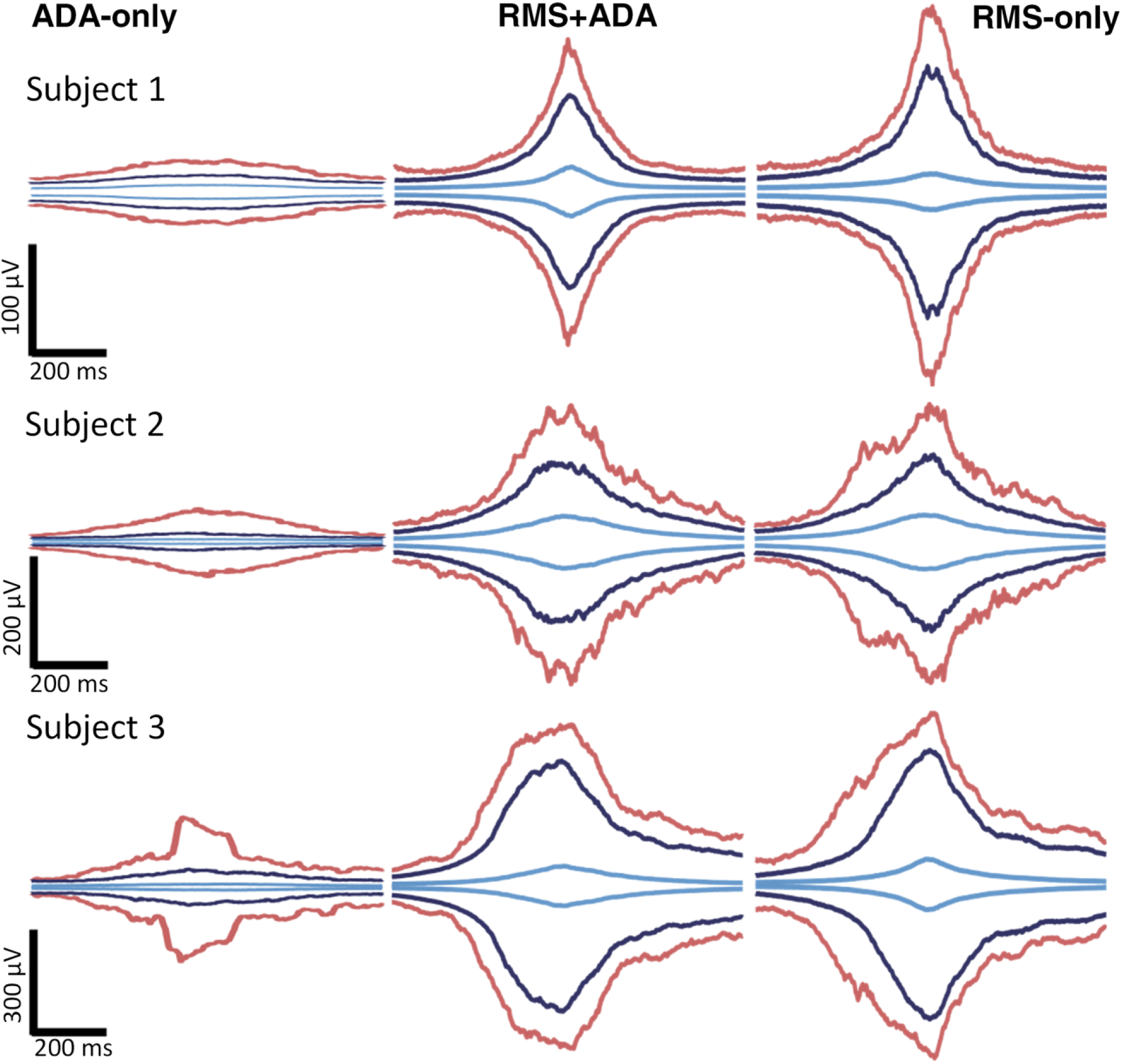
Amplitude envelopes of detected events separated into ADA-only (left panel), RMS+ADA (center panel), and RMS-only (right panel) from three representative subjects. The maximum, 95^th^ percentile, and median amplitude envelope are represented by red, blue, and light blue lines, respectively.

### 3.2 Characteristics of aHFA and cHFO

Because the rate (average number of HFOs per minute) has been almost exclusively used as a biomarker of the SOZ, we first measured the aHFA rate and cHFO rate in individual subjects, as well as the total rates when all events were pooled together (Figure 5A). Our results for rate using the RMS detector were consistent with previous studies: the rate of cHFO in the SOZ was 7.8±5.0 per minute, which was significantly higher than the rate in the nSOZ, 1.8±2.9 per minute. The individual results for all 11 patients also exhibited significant differences in cHFO rate between SOZ and nSOZ. The results for rate using ADA were less consistent. The average rate of aHFA was significantly different between the SOZ (3.5±2.9 per minute) and nSOZ (3.9±2.2 per minute), with the nSOZ having a higher rate. However, only six out of eleven patients had significantly different rates of aHFA in SOZ and nSOZ. In five out of these six patients, higher rate of aHFA was observed in nSOZ. Therefore, the rate of aHFA does not provide reliable separation between SOZ and nSOZ channels.

**Figure 5:**
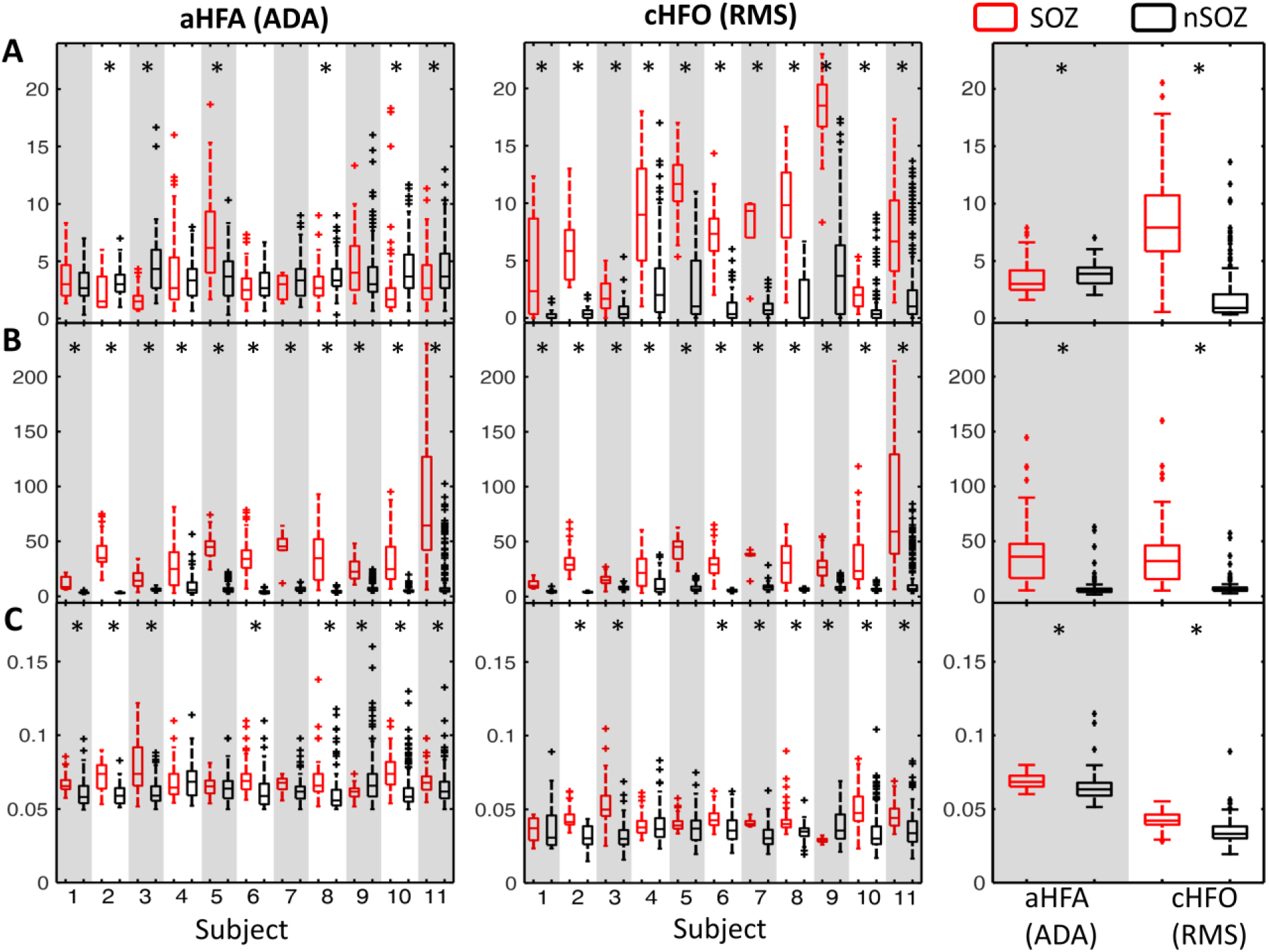
Characteristics of detected events separated into SOZ (red) and nSOZ channels (black). Boxplots show the (A) rate, (B) amplitude, and (C) duration of detected events for individual subjects (aHFA in left column and cHFO in middle column) and overall characteristics of detected events when all segments are pooled together (right column). * p-value < 0.0045, Wilcoxon rank sum test with Bonferroni correction for 11 subjects.

In contrast to the rate, the event amplitude showed robust differences between SOZ and nSOZ channels using both ADA and the RMS detector (Figure 5B). The mean amplitudes of cHFO (38.0±33.7 µV) and aHFA (40.1±32.5 µV) in SOZ were significantly higher than in nSOZ (8.8±9.1 and 7.1±9.1 µV, respectively). These differences were statistically significant for all 11 individual subjects using both detection methods. Moreover, the amplitudes were consistent across subjects, such that a single, common threshold of approximately 15 µV can approximately separate SOZ and nSOZ across all subjects. This is not true for the rate, which requires selection of a patient-specific threshold to separate SOZ and nSOZ channels. The robustness and consistency of these results suggests that the amplitude of aHFA and cHFO may be potential biomarkers of the SOZ.

The average duration of detected events within SOZ channels was significantly longer than the duration in nSOZ channels for both detection schemes (Figure 5C). This difference was statistically significant for 8 subjects using ADA and 9 subjects using the RMS detector, with events in the SOZ generally having a longer duration. However, for individual subjects, the duration of cHFOs provided smaller separation between SOZ and nSOZ channels than event amplitude or rate.

Finally, we calculated the CV to assess the consistency of the measurements across segments of data (Figure 6). A low CV value indicates a consistent measurement, where the SD is lower than the mean. The amplitude CV was significantly lower than the rate CV when calculated using the same detector and channels, suggesting that the estimates of amplitude are more stable over time for both detection schemes. The CV for amplitude in the SOZ was higher than in nSOZ, but it was still generally lower than the CV values for rate.

**Figure 6:**
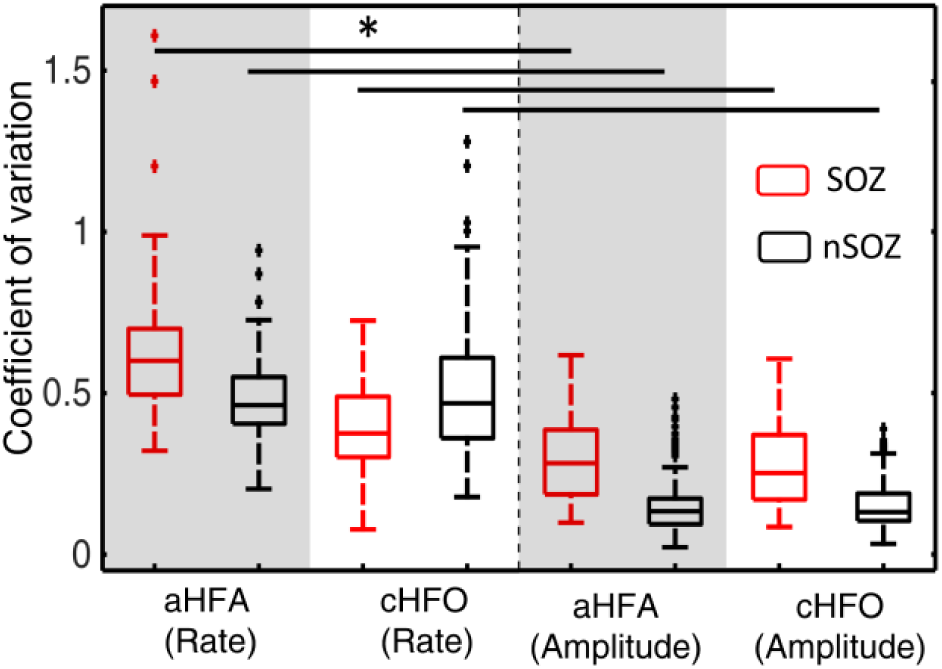
Boxplots of CV for the rate and amplitude of detected events, separated into SOZ (red) and nSOZ (black). * p-value < 0.0125, Wilcoxon rank sum test.

## 4 Discussion

Here, we have presented a novel algorithm for detection of anomalous high frequency events in electrophysiological data and applied it to human iEEG. ADA is unsupervised and does not require complex optimization procedures or prior assumptions about the shape or amplitude of the events. While the rate of aHFA was not consistently different between SOZ and nSOZ channels, the amplitude of aHFA provided reliable and robust separation between SOZ and nSOZ channels, suggesting this parameter as a possible biomarker of the SOZ. We found significant overlap between aHFA and cHFOs, indicating that ADA is sensitive to traditionally-defined HFOs. However, we also found that ADA identifies additional events that would not exceed the energy threshold of standard algorithms.

Robust identification of clinically-relevant HFOs has remained a challenge due to the lack of a physiological definition of these events and the limitations of current detection methods. Because the mechanism underlying HFOs is not fully understood, detection is typically guided by the empirical definition which requires selection of an optimum energy threshold to separate events from the background. This is not a trivial task because the shape and amplitude of the HFO waveform can vary depending on the distance between the electrode and the neural generator^39^ and the characteristics of the background activity vary over time. Therefore, a rigid template of shape and amplitude is likely to be insufficient for HFO detection. In addition, current automated algorithms require complex optimization procedures to maximize accuracy,^17^ and it is common for them to suffer from a high number of falsely detected events.^17,18,23^ Visual identification by expert reviewers has been widely used for detecting HFOs in both scalp and iEEG recordings,^40,41^ as human reviewers can simultaneously adapt to changes in the background activity and perform artifact rejection. However, it is highly time-consuming and has poor agreement between reviewers,^19^ which reduces the generalizability of the results. ADA addresses these challenges, as it does not require human input and can therefore perform unbiased detection and estimation of event characteristics.

In the present study, the default parameters for ADA were configured to detect anomalous events at frequencies greater than 80 Hz. The small window was used to downsample the data to reduce the calculation time;^42^ the approximate detection time for three minutes of data from a single channel of iEEG was ∼4-5 minutes using a desktop PC (CPU: i7-4790k). With parallel computing or optimization of the algorithm, it may be possible for the size of this window to be reduced or to exclude this step from the procedure. The size of the 1.5 ms small window is equivalent to downsampling of the signal to 666 Hz, which means that our algorithm is primarily detecting events in the ripple frequency band (80-250 Hz). However, the algorithm could be configured for other frequency bands as well, simply by changing the sizes of the small and large windows. The large sliding window was chosen to match the approximate duration of HFOs reported in prior studies.^21,26,43^ Because the algorithm contains a step to join overlapping large windows in the anomaly group into single events, a range of large window sizes can be used without affecting the results. In the clustering and classification process, the maximum number of clusters was seven; however, this number can be altered, as long as it is higher than the expected number of anomalous patterns. We tested the algorithm with a range of seven to thirteen clusters, and there was no noticeable difference in the results in this range because the background cluster is always several orders of magnitude larger than any other cluster. Therefore, we chose to use seven clusters, as it reduced the processing time.

We found that aHFA had a longer duration than cHFO, but there were significant differences between the two detection algorithms that contributed to this result. The aHFA had a minimum duration of 50 ms, corresponding to the size of the large sliding window, while the cHFO duration was measured as the length of time that the RMS amplitude exceeded the energy threshold. This impacted the measurement of amplitude, as well. The same event detected with ADA may have a lower amplitude than when it is detected with the RMS detector because the average amplitude will include some background activity at the edges of the window.

Conventionally, the rate of cHFO has been used as a biomarker of the SOZ in studies of high frequency activity related to epilepsy.^2–9^ Here, we found that the rate of aHFA was less predictive of the SOZ than the rate of cHFO detected using a strict energy threshold, as ADA detected many low amplitude events which were not detected by the RMS detector. Every additional detected event raises the rate by one event per minute, similar to the effect of false positive detections when using conventional automated algorithms. This can drastically change the relative rates in SOZ and nSOZ channels (especially for ADA, which more frequently detects low amplitude events in nSOZ) and can therefore alter the prediction of SOZ location. This is supported by the fact that the CV of the rate was higher than for amplitude or duration, indicating a higher degree of variability across segments of iEEG.

In contrast to the rate, the aHFA and cHFO amplitudes were significantly higher in the SOZ compared to the nSOZ, with relatively consistent values across patients. Several other studies have suggested that HFO amplitude differs between SOZ and nSOZ channels. One study has shown amplitude to be significantly higher in SOZ compared to non-SOZ electrodes (regardless of contact location) during interictal, preictal, and ictal periods,^11^ but others have shown that this difference was not significant during nonictal periods.^44^ Furthermore, pathological HFOs have been reported to demonstrate higher amplitude than physiologically-induced HFOs.^26^ However, because the reported differences were generally small, amplitude has not received much attention as a possible biomarker of the SOZ.

Here, we found that the amplitude of cHFO, amplitude of aHFA, and the rate of cHFO provided robust separation between SOZ and nSOZ in all 11 patients. However, the amplitude exhibited less variability over time and more consistency across patients. This suggests that amplitude may be another promising candidate for a biomarker of the SOZ. Further validation with a larger cohort of patients, more comprehensive inclusion of iEEG electrodes, and comparison to surgical outcome are needed to explore this hypothesis and will be the subject of future investigations.

## Acknowledgements

This research was financially supported by Royal Thai Government Fellowship awarded to K. Charupanit

